# Climate stress and its impact on livestock health, farming livelihoods and antibiotic use in Karnataka, India

**DOI:** 10.1101/2022.06.10.495626

**Authors:** Adam Eskdale, Mahmoud El Tholth, Jonathan D. Paul, Jayant Desphande, Jennifer Cole

## Abstract

Understanding the impact of climate change on livestock health is critical to safeguarding global food supplies and economies. Informed by ethnographic research with Indian farmers, veterinarians, and poultry industry representatives, we evidence that both precipitation and vapour pressure are key climate variables that relate to outbreaks of haemorrhagic septicaemia (HS), anthrax (AX), and black quarter (BQ) across the Indian state of Karnataka. We also identify temperature and maximum temperature to be negatively correlated with the same diseases, indicating that a cooling (but still hot) climate with wetter, humid conditions is a prime risk factor for future outbreaks. Principal component analyses have revealed the SW India monsoon and winter periods to be the most strongly correlated with HS, AX and BQ outbreaks. We identify vapour pressure, a proxy for humidity, as having a positive relationship with these specific livestock diseases. The negative relationship between temperature and these diseases, combined with the positive correlation with rainfall and humidity, allow us to classify climate-associated risk using a combination of gridded meteorological time series and epidemiological outbreak data covering the same region and timespan of 1987–2020.

Risk maps were constructed following concerns over the growing impact of climate pressures raised by farmers during ethnographic study. Informed by their insights, we used current climate data and future climate projections as a risk classification tool to assess how disease risk varies in Karnataka in the present and possible future scenarios. Despite a relatively limited epidemiological dataset, clear relationships between precipitation, vapour pressure, and temperature with HS, AX and BQ, along with outbreak high-risk zones were defined. This methodology can be replicated to investigate other diseases (including in humans and plants) and other regions, irrespective of scale, as long as the climate and epidemiological data cover similar time periods. This evidence highlights the need for greater consideration of climate change in One Health research and policy and puts forward a case for, we argue, greater alignment between UNFCCC and One Health policy, for example, within the Tripartite Agreement (between OIE, FOA and WHO) on antimicrobial resistance as disease risk cannot be considered independent of climate change.

**One Health Impact Statement:** This research aims to investigate the relationship between factors related to climate (surface temperatures, rainfall, humidity) and outbreaks of livestock-related bacterial diseases. This is especially relevant to the One Health approach as it attempts to integrate findings between not only the science of disease but also the science of climate change as a driver of disease, and address problems that could arise within the public and private sectors (local farming, livestock health, government policy etc.). Providing spatial context to climate-associated disease risk across the Indian state of Karnataka will benefit local farmers that may already be, or transitioning to, more intensive livestock farming along with policy makers and private sector companies who are planning for future investments. This transdisciplinary approach springboards from ethnographic observations of famers’ lived experiences of challenges to their livelihoods and facilitates the use of climate datasets that may not have been primarily collected for or used by disease-related studies to map long-term epidemiological risk. This demonstrates the pragmatic impact that such transdisciplinary projects can have by providing interpretations of observed risks to animal health (highlighted by social scientists during engagement with practitioner communities) that Earth Scientists were then able to quantify, proving links that would be otherwise not have been evidenced. Using disease data sourced from local institutions, including Government of India facilitates as well academic research laboratories, can plan the application of pragmatic solutions to local farmers who are primarily impacted by the findings of the research.

## 1 Introduction

India is currently ranked as the nation that was fourth most-affected by climate change between 1996 and 2015 (Kreft, Eckstein and Melchior, 2017). There is particular variance in the climate changes between the northern and southern parts of the country, with both becoming increasingly warmer over historic meteorological trends (Dash and Hunt, 2007). At the base of the Himalayas, surface temperatures extremes are increasing: hotter in the summer months and colder in the winter (Dash and Hunt, 2007; Dash *et al*., 2007; Sanjay *et al*., 2020). Monsoon precipitation increasingly fluctuates, predicted to increase across India into the near future; monsoon rains also tend to start earlier as a result of anthropogenic aerosols (Bollasina, Ming and Ramaswamy, 2013; Kulkarni *et al*., 2020). This temporal shift in precipitation along with increased extreme values is linked to increasing susceptibility to droughts and associated hazards with such environments in some areas (wildfires, groundwater fluctuation), and in others with significant rise in wet-bulb temperatures (i.e., humidity; Sinha and De, 2003; Prabhakar and Shaw, 2008; Sahu, Sett and Kjellstrom, 2013; Mujumdar *et al*., 2020). These interlinked climatic factors will essentially make livelihoods of local populations more insecure and precarious in the coming decades, one such consequence being the potential for increased risk in livestock bacterial, viral, and parasitic disease. In turn, this threatens to increase the use of antibiotics to combat disease risk, exacerbating the already dangerously high use of antibiotics in the Indian livestock sector that drives antibiotic resistance (Mutua *et al*., 2020).

Increasing climate variability will continue to have profound impacts on global health and food supplies (Gregory, Ingram and Brklacich, 2005; Shukla *et al*., 2019). It is becoming increasingly well documented that changes in land temperature, rainfall and diurnal temperature ranges can have important relationships with disease (Rohr *et al*., 2011; Messina, 2019; Messina *et al*., 2019; Vinke *et al*., 2020). Further understanding of these relationships between climate changes and disease patterns is critical to ensuring future societal health and food security, as well as animal health and farming livelihoods, is maintained.

Climate change affects the health of humans, animals and plants directly, through heat and cold that affect body temperature and plant growth, via extreme weather that causes floods or droughts, and through indirect impacts on food production, air quality and other environmental factors (Cramer *et al*., 2018). India is at particular risk of climate change-related health impacts (Majra and Gur, 2009; Singh and Dhiman, 2012), which are predicted to increase in the coming decades, as average temperatures could increase by as much as 2°C by 2050 (World Bank Group, 2022). Of particular concern to India is the impact of climate change on key food production industries such as the poultry sector (Pawar *et al*., 2016), which threatens the loss of economic development opportunities as well as food security. In addition to the direct impact of climate change on the poultry industry, climate stress has also been shown to exacerbate other challenges, such as the misuse of antibiotics in poultry production to treat symptoms of heat stress that mirror those of bacterial infection, reducing the efficacy of antibiotics. Where this happens at the same time that infections they are needed to treat are likely to increase due to heat stress and other climate-impacted factors such as harder water from deeper borewells reducing the efficacy of cleaning products, and increased ranges of disease-carrying parasites (Cole and Desphande, 2019), action plans to control disease are unlikely to be effective they do not fully consider the impact of and challenges raised by climate change.

We began by attempting to understand drivers of antibiotic use and misuse in the Indian livestock sector. During ethnographic work undertaken in peri-urban areas within a 25 km radius of Bengaluru, Karnataka in southern India during 2018–19, farmers consistently spoke of challenges to their livelihoods from climate change (Cole and Desphande, 2019; Greru *et al*., 2022), while veterinarians pointed to misuse of antibiotics to treat symptoms they suspected were caused or exacerbated by heat stress rather than bacterial infection (Cole and Desphande, 2019). This suggested a direct relationship between climate stress and disease which required further examination, especially as climate stress is increasingly pushing farmers from crop raising to less water-intensive poultry/livestock production, which could also exacerbate climate change issues in the long term due to land use changes and increased energy requirements (McMichael *et al*., 2007). As the Global North attempts to shift to a more plant-based diet to combat carbon emissions and environment damage caused by livestock farming (Willett *et al*., 2019), combined with the large vegan population (Wright, 2021), the Indian farming sector could find itself left behind if the impact of climate change on its practices is not fully understood and planned for. In exploring the reasons for inappropriate use of antibiotics in livestock health and veterinary practices, it is important both to understand the lived experience of farmers who are seeing the challenges first hand (Badstue *et al*., 2018), and to be willing to take a systems approach that seeks to understand more complex interplays of drivers and usage pathways. However, at present, dialogue between ecosystem science, climate change and public health on topics such as antimicrobial resistance could be improved (Iossa and White, 2021). Our aim is to present a case study that helps to links these two disciplines (climate science and One Health) and show to those working in each field the potential benefits of working more closely together.

### 1.1 Aims and Objectives

Recognising how climate changes may relate to livestock diseases is essential to predicting future outbreaks and planning future farming policies. We sought to listen to the concerns of the farmers who were participants in the ethnographic research and leverage their experience to help us anticipate the impacts climate change may have on Indian livestock farming in future, by exploiting existing climate data to identify the most acutely affected regions. Secondly, we aim to develop a better understanding of how climate impacts poultry and livestock health; and finally, we develop a methodology through which other researchers and practitioners – including farmers, veterinarians, industry representatives and policymakers – can better understand climate risks to livestock.

The purpose of this paper is therefore to assess the usefulness of integrating lived experience of farmers, veterinarians, and poultry industry representatives with meteorological and epidemiological datasets to identify and evaluate relationships between climate and livestock diseases (here, specifically bacterial disease).

The key objectives are to:

1. Identify climate changes that relate to livestock bacterial diseases in Karnataka.
2. Create risk assessment maps that provide spatial context to these relationships, identifying higher and lower risk areas based on the contributions of each variable.
3. Project these findings onto future climate data, constructing prediction maps for areas of higher risk and lower risk of bacterial disease based on climate observables.
4. Develop a methodology for calculating risk that can be automated and embedded into a user-friendly platform that can be used by practitioners and policymakers, not necessarily climate science experts.

The remainder of this paper is structured as follows. In Section 2 the methodology for this study is outlined, describing the research process and each dataset component used. In Section 3 the results are presented, identifying the relationships from these data, and risk maps are constructed. Finally, Section 4 discusses the overall impacts of these findings and how the risk maps can be applied by both farmers and government, before presenting brief conclusions and a future outlook.

## 2 Methods

### 2.1 Study Area: Karnataka

Karnataka is one of five Indian states that produce more than 60% of broiler chickens and eggs (the others being Andrah Pradesh, Maharashtra, Punjab and West Bengal: Dubey *et al*., 2021). The city of Bengaluru (Fig. 1) hosts a number of animal science and veterinary institutes, such as the National Institute of Veterinary Epidemiology and Disease Informatics (NIVEDI), the National Dairy Research Institute (NDRI), the Central Poultry Development Organization (CPDO) and the Karnataka Veterinary Animal and Fisheries Science University (KVAFSU). It is thus an important centre of animal production and research in India. The research team contained and interacted with staff from many of these institutes as part of the projects DARPI and NEOSTAR, funded under the same programme (Shaju, 2017) and visited several dozen farms during their research (e.g., Cole and Desphande, 2019; Greru *et al*., 2022) conducting ethnographic observations and key stakeholder interviews with farmers, veterinarians, members of the poultry industry and policymakers.

**Figure 1:**
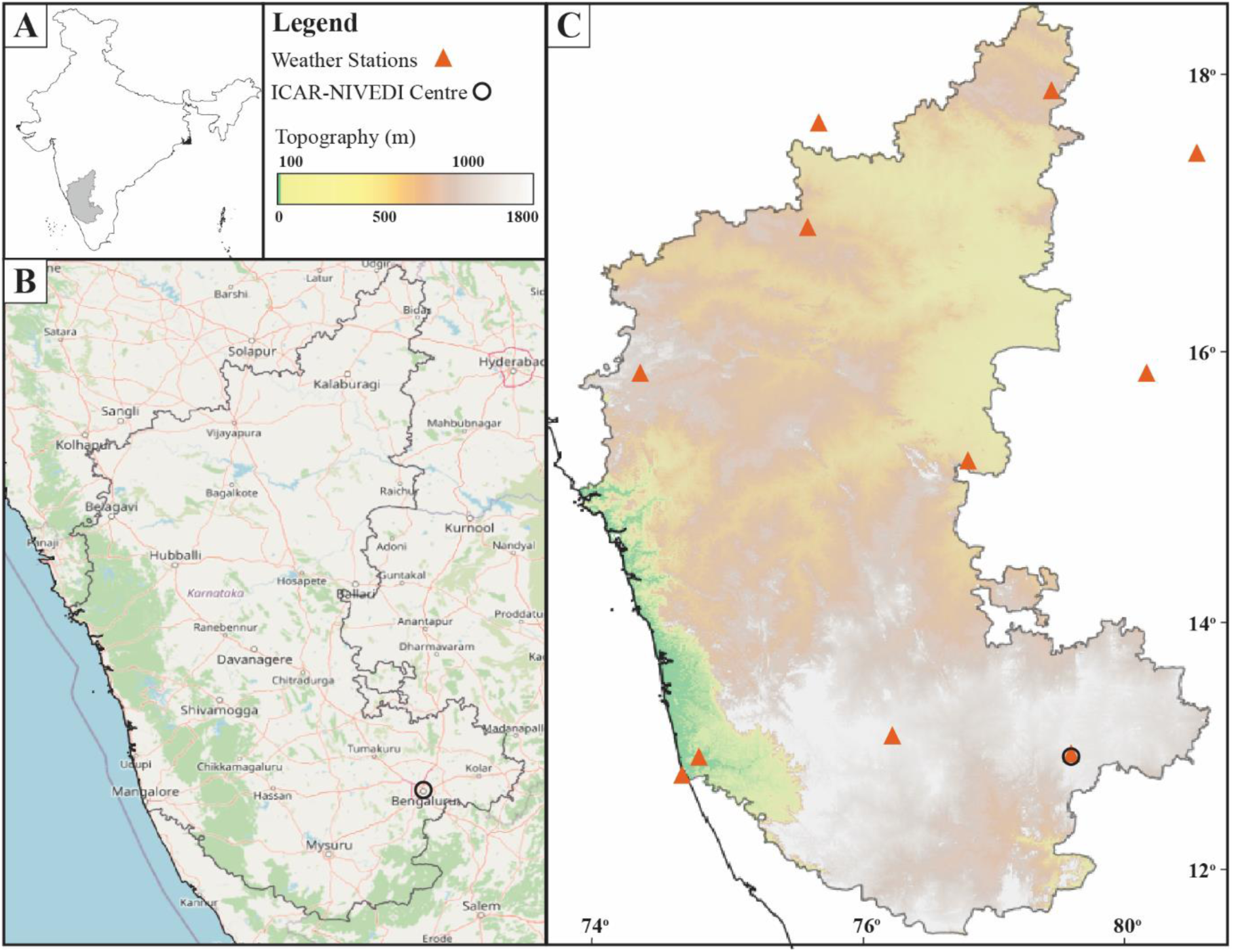
Location (A,B) and topographic (C) map for the state of Karnataka, India.

Karnataka has been particularly prone to climate stress in recent years, especially from drought and rising temperatures (Srinivasareddy *et al*., 2019; Lokesh *et al*., 2020), to which the key stakeholders often made reference. Local average temperatures are predicted to rise by 1.8–3.3°C in Karnataka by 2030 with respect to the baseline period 1961–90 (Murari *et al*., 2018). The links between climate change, increasingly precarious farming livelihoods and thus changes to farming practices (some of which increased the use of antibiotics in the farming sector) were raised by farmers interviewed during ethnographic stages of the DARPI and NEOSTAR projects. This prompted detailed investigation of potential links between climate variables and disease in Indian livestock production and health, which had not originally been intended to be part of either project. We leveraged the lived experience of the stakeholders to identify the ability of the region of India in which the DARPI project was undertaken (sufficiently robust epidemiological data relating to the region in which NEOSTAR was undertaken was not available) to sustain livestock production in future and to predict which other regions may become less or more able to sustain livestock production or may require additional mitigations to enable current activity to continue. Exploring this understanding will help farmers, livestock sector representatives and policymakers to plan future operations and expansion with these climate factors in mind.

### 2.2 Disease database

Bacterial disease outbreak data for Karnataka were sourced from the NADRES (National Animal Disease Referral Expert System) database (NADRES, 2017), maintained by NIVEDI; data are input to the database by farmers who self-report livestock diseases. These data are not followed up for subsequent investigation and we acknowledge the limitations of this. Data were extracted from the database and collated manually; since the earliest available data for Karnataka is 1987, we focus on the subsequent 33-year period of 1987–2020. Data for a total of 15 different livestock-associated diseases are available in the database: Anthrax, Babesiosis, Black Quarter, Bluetongue, Contagious caprine pleuro pneumonia, Enterotoxaemia, Fascioliasis, Foot and Mouth disease, Haemorrhagic Septicaemia, Peste des petits ruminants, Rabies, Sheep and Goat pox, Swine fever, Theileriosis, and Trypanosomiasis. Only bacterial diseases were selected to relate to the research interest with the misuse of antibiotics, therefore restricting our selection to four diseases – Anthrax, Black Quarter, Enterotoxaemia, and Haemorrhagic Septicaemia. The disease data are not unique to a particular species, but cover five major livestock species: buffalo, cattle, sheep, pigs, and goats. Hereafter we refer to these collective species as ‘livestock’. The interpretations from these data can therefore not be disaggregated to a specific species as the data does not allow this; however, we believe that it remains useful.

### 2.3 Climate Data

Climate data used in this study were sourced from the Climatic Research Unit (CRU) TS 4.5 dataset (Harris *et al*., 2020). Data were downloaded and cleaned before statistical and correlation analysis was undertaken. The CRU dataset was used as it contains climate data from 1901–2021 including all climate variables that were of interest to this study (i.e., temperature, maximum temperature, vapour pressure, precipitation, diurnal temperature range). In addition, the CRU dataset is easily accessible through a WPS server and the Google Earth interface and has a high spatial resolution (0.5 x 0.5°). These climate variables were selected as the variance is easily measurable, they are the main parameters that fluctuate during seasonal shifts, and also have established relationships with other diseases (Cheng *et al*., 2014; Escobar *et al*., 2017; Ezenwa *et al*., 2020). Vapour pressure was especially important to involve as it is a proxy for humidity (Shamshiri *et al*., 2018), and there are very few studies that define relationships with this variable and livestock bacterial disease.

### 2.4 Climate and Disease Data Modelling

Climate and epidemiological modelling were conducted in a three-stage process.

1. Disease and climate data were collected and then separated into annual, monthly, and seasonal groups for easier comparison. Seasons were defined using the generally accepted months for each main period in South-West India: Winter (Jan-Feb), Summer (March-May), Monsoon (June - Sept), and Post-Monsoon (Oct-Dec). Further data, such as climate anomalies, were generated for later use. Climate data for each state (treated as a whole) were collated by combining all 0.5 x 0.5° grid box data that cover the states into one monthly average.
2. The relationships between the climate and disease data were investigated using multiple correlative statistical analysis techniques, using both Pearson’s and Spearman’s Rank, followed by a principal component analysis.
3. Climate data and climate anomalies, disease risk and locality were mapped. Ranges for disease risk were first categorised, based on the interpretations from the statistical analysis.

### 2.5 Predicting Bacterial Disease Outbreaks

Risk categories were assigned based on percentage quantiles using the relative percentage difference (RPD) values for each climate variable in each season from the total period (1987-2020) mean. For example, the ‘very high risk’ category for precipitation is any grid box that has RPD values in the top 0.8 percentile deviating from the 1987–2020 average. This classification system was conducted per climate variable, assigning a numerical value of 1 to 5 depending on the outcome (1 being the lowest risk, 5 being the highest). It was also repeated per season for each variable, facilitating the impact of seasonal variation in the risk assessment. The sum of the overall risk for each climate variable was then used to classify a total risk number, again using percentiles. For example, those areas that were in the top 0.8 percentile using the sum of each climate variables risk are then classified as ‘very high risk’ areas overall.

## 3 Results

### 3.1 Bacterial Disease Outbreaks and Climate Variable Trends, 1987– 2020

By comparing several decades of climate data to a similar period of bacterial disease outbreak data, we were able to define correlative relationships. First, we matched qualitatively important maxima and minima.

#### 3.1.1 Long-term Climate Trends

On average across the state of Karnataka, four of the five climate variables (precipitation, vapour pressure, maximum temperature, and surface temperature) have been increasing from 1987 to 2020 (Fig. 2). Precipitation and vapour pressure fluctuate significantly between specific years, but trend positively overall especially when considering the five-year running average trendline. Surface temperature and maximum temperature also increase, but fluctuate less than precipitation and vapour pressure, remaining relatively stable despite a few anomalous years (e.g., 1998, 2010). Diurnal temperature range fluctuated between 1987– 2002, after which the records become stable at ∼10.85^°^C. This stability is a clear artefact of the dataset rather than reflecting true values, as the data remains at this value for 18 years which is unlikely. Due to this distortion through the majority of this selected time period, diurnal temperature range is omitted from Fig. 2.

**Figure 2:**
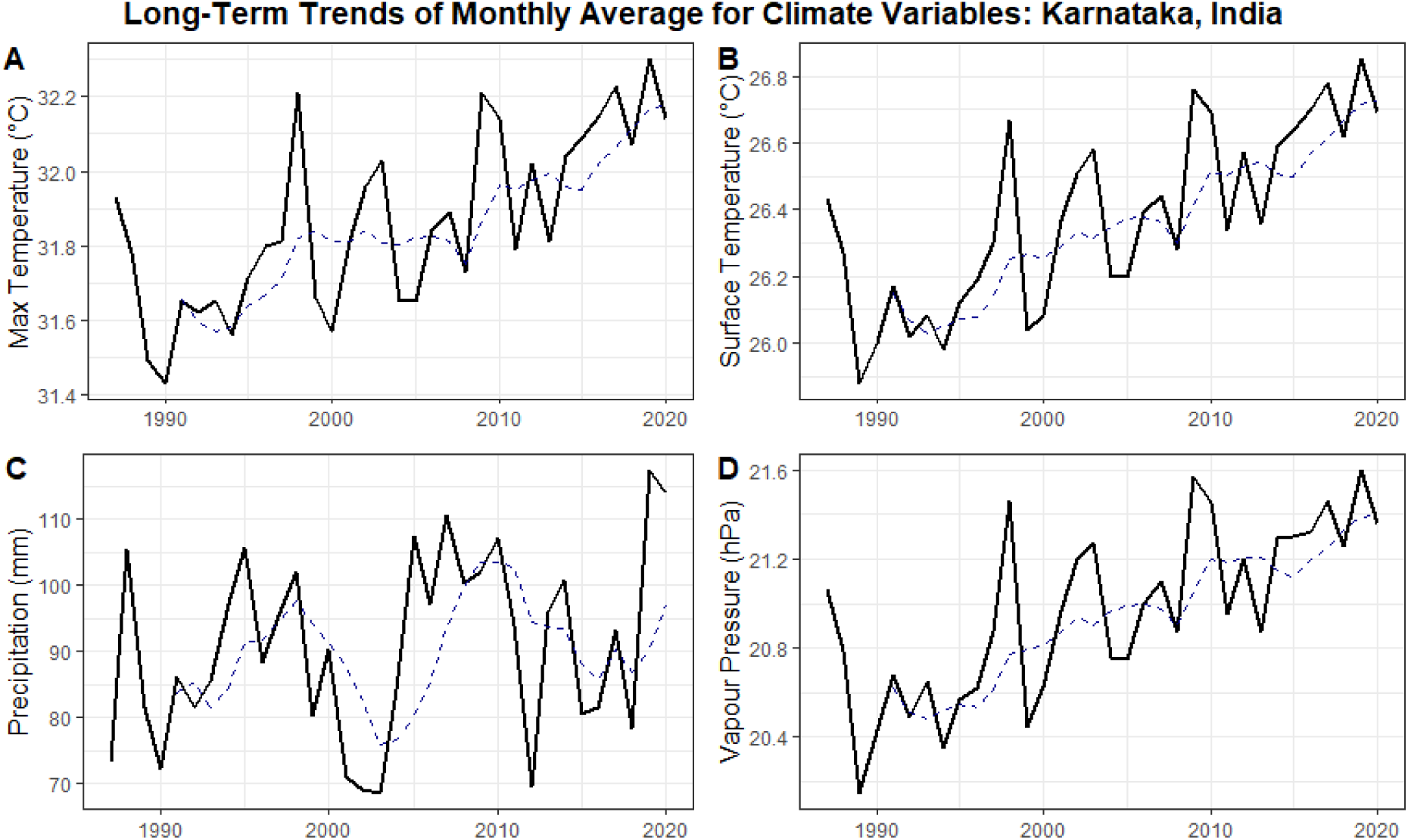
Monthly averages of climate observables, 1987–2020, across the state of Karnataka, India. (A) Maximum surface temperature; (B) Surface temperature; (C) Precipitation; (D) Vapour pressure. Dotted lines = five-year moving average. Data from Harris et al., (2020).

Such nuances in climate data are relevant, as many livestock animals homeostatically regulate core body temperature within a narrow range (Biswal *et al*., 2022). Dairy cattle, for example, need to maintain a constant body temperature of around 38.8±0.5°C (Ohnstad, 2008). A high ambient temperature above this thermal comfort zone, caused by climate stress, will trigger series of neuroendocrine modulations that are detrimental to the animals’ welfare and productivity. In broiler chickens raised for meat production, heat stress (HS) will reduce feed consumption, growth rate, feed digestion and efficiency, immunity, survival rate and overall welfare (Abioja and Abiona, 2021). Dairy cows will experience lethargy, declines in feed intake and thus milk production, reduced fertility and an increase in susceptibility to mastitis (Dahl, 2018). Climate change could also impact disease patterns via changing migratory routes for wild birds or other disease vectors: highly pathogenic Avian Influenza is an example that could spread wider, while diseases such as blue tongue have also increased in geographic distribution in recent years due to climate variation (Mayo *et al*., 2014). New strains of livestock diseases could also emerge via mutations influenced by climate change (Gale *et al*., 2009). It has been suggested that seasonal fluctuations not only influence the distribution of current bacteria, viruses, parasites and their vectors but also the emergence of new diseases (Ahaotu *et al*., 2019).

#### 3.1.2 Long-term Bacterial Disease Trends

Publicly available, accurate to high granularity (individual village / town), and temporally complete bacterial disease data for India are quite limited, with one of the main sources being the NADRES v2 database. Data are available for the number of outbreaks per disease from 1987–2020 through an online GIS portal, although data availability can fluctuate, most likely due to the reliance on data providence from farmers alone. For this study, disease data were available for Karnataka over 1987–2020 (Fig. 3).

**Figure 3:**
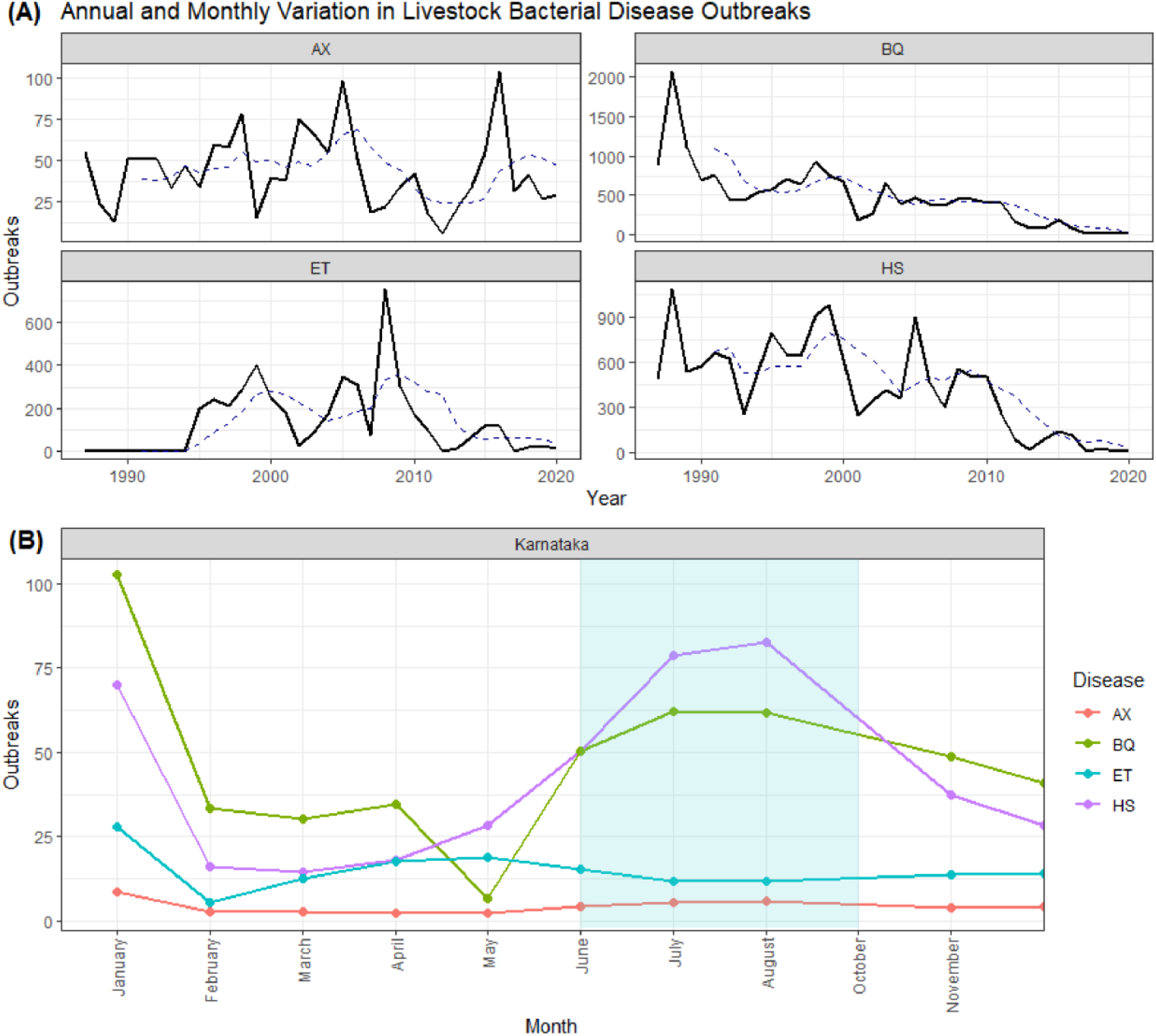
1987–2020 trends for bacterial disease outbreaks in livestock in Karnataka. (A) Haemorrhagic Septicaemia (HS), Anthrax (AX), Black Quarter (BQ), and Enterotoxaemia (ET); dashed line = five-year moving average. (B) Monthly averages of disease outbreaks, 1987–2020, with monsoon months shaded in blue. Data sourced from NADRES database.

In Karnataka, HS outbreaks decreased over the study period (Fig. 3A). Black Quarter also decreases with less variability, while the annual number of outbreaks of Anthrax varies greatly, making it harder to infer longer-term trends. Enterotoxaemia outbreaks increase, with significant peaks in 1999, 2005 and 2008.

Haemorrhagic septicaemia is most prevalent during the SW monsoon months (June to September), and predominantly low during the summer months (March to May: Fig 3B). Anthrax outbreaks remain relatively low throughout the year compared to the other diseases; however, there is still a subtle increase during the SW monsoon months, and a relatively low through the summer. Black Quarter also peaks during these months; BQ outbreaks are lowest during May. During the summer months ET has a shallow, wider peak in Karnataka before then decreasing to a lower, stable level throughout the remainder of the year at around 17 outbreaks per month. All of the bacterial diseases have peaks in January followed by a sharp decrease, indicating distinct change from the winter months into the summer (Fig. 3B).

### 3.2 Seasonality of Disease Outbreaks vs Climate Variability

Although identifying average trends throughput the year is useful, comparing averaged seasonal climate data with disease data for the same period allows for a clearer interpretation of differences in outbreaks per season: first, by comparing peaks between long-term trends, then by identifying potentially more subtle relationships.

#### 3.2.1 Peak-to-Peak Correlations

Bacterial disease outbreaks noticeably vary in Karnataka (Fig. 4). The long-term trends for BQ and HS indicate they decrease over time, though both fluctuate annually; the rate of this decrease varies per season with more notable definition within the monsoon. The outbreaks of AX remain relatively lower and more stable, increasing slightly in 2005. Outbreaks of ET are higher than AX and fluctuate slightly more but are still relatively stable compared to HS and BQ.

**Figure 4:**
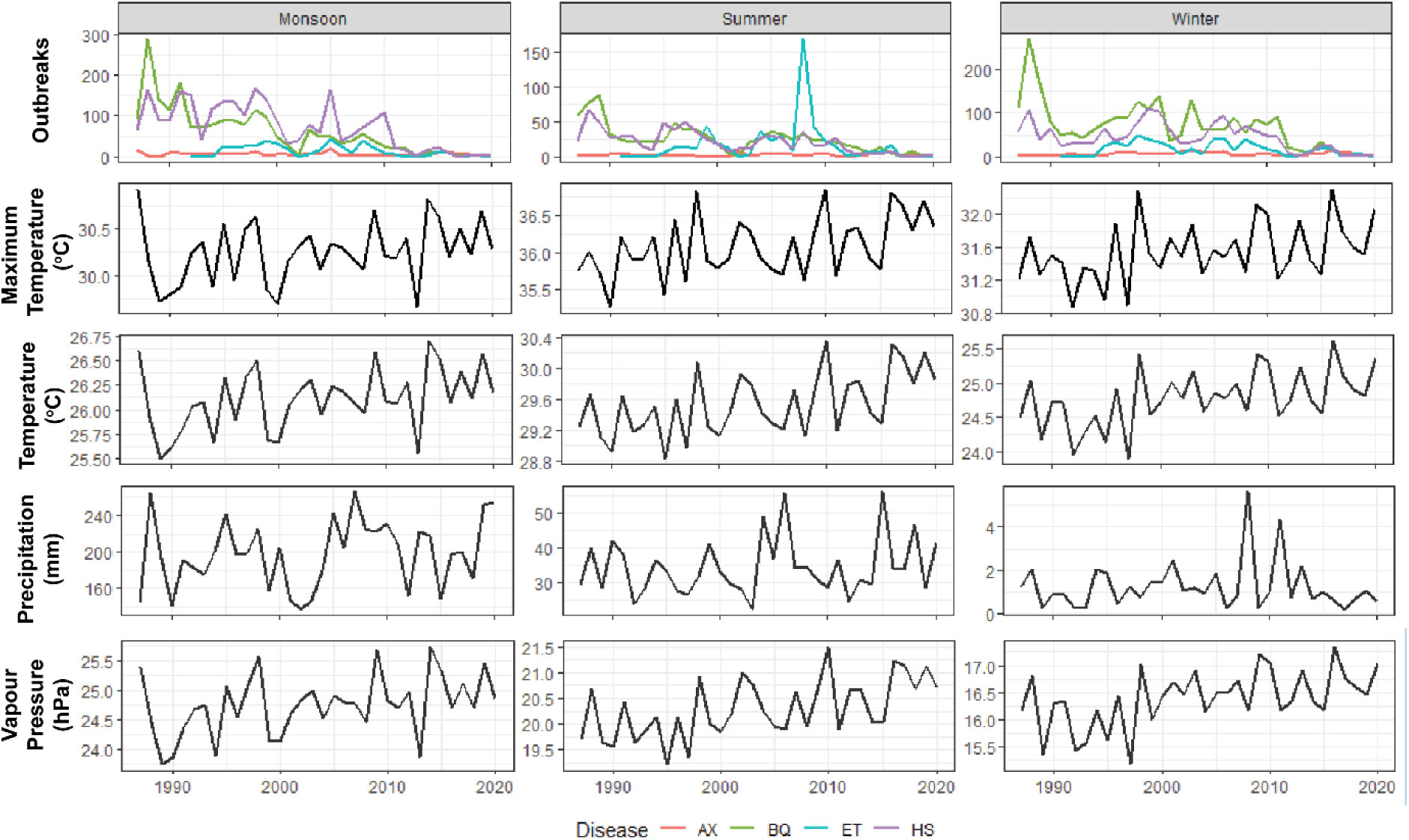
Long-term bacterial disease and climate trends for the monsoon, summer, and winter periods over 1987– 2020. First row: Karnataka bacterial disease outbreaks; Second to fifth rows: monthly averages of maximum temperature, surface temperature, precipitation, and vapour pressure, respectively.

Monsoon temperatures in Karnataka are increasing, despite year-on-year fluctuation (Fig. 4). Temperature and maximum temperature correlate negatively with some bacterial diseases but positively with others. Exemplar years include 1997 and 2005 where a temperature spike coincides with a spike in disease, and in 1992, where a decrease in temperature matches a spike in HS and BQ. Precipitation remains relatively stable on average for the monsoon period, and correlates with peaks in bacterial disease, with spikes matching in the years 1990, 1995, and 2010. These spikes are clearer between precipitation and HS, BQ and ET, while it is harder to discern AX peaks. The one clear peak in 2005 of all four bacterial diseases matches a decrease in precipitation, indicating there may also be mixed and complex relationship alike with temperature. Vapour pressure remains relatively stable throughout the period, with notable peaks in 1997 and 2003 aligning with peaks in HS, BQ and ET indicating a possible positive relationship (Fig. 4). Finally, diurnal temperature range was omitted from the peak-to-peak analysis as the data appears to remain the same value repeatedly for 11 years - most likely artificial and not reflecting natural fluctuations and is therefore impossible to interpret any correlation visually.

Through the summer HS, AX and BQ all decrease over the 33-year period while ET increases modestly (Fig. 4). All diseases fluctuate year by year, with distinct peaks of HS and BQ in 1988–89, and a major peak of ET in 2009. The temporal pattern of extrema is similar in both the precipitation and ET datasets, for example matching peaks seen in 2000, 2004, and 2006 with the exception of 2016, where the patterns are the opposite (Fig. 4). Overall maximum temperature, surface temperature and vapour pressure have mixed correlations with the bacterial diseases.

Average winter values for all the bacterial diseases in Karnataka decrease over 1987–2020. Throughout the winter HS outbreak peaks correlate positively with small peaks in vapour pressure (Fig. 4: 1988, 2016), but also correlate negatively in some years (Fig. 4: 1995, 2008). Black Quarter peaks correlate positively with vapour pressure, average temperature, and maximum temperatures (Fig. 4: 1999, 2003) but also negatively (2011). Outbreaks of BQ correlate both positively and negatively with precipitation (Fig. 4: 2011, 2003). Precipitation peaks match ET outbreaks in 2008, but this is not a consistent pattern. Overall AX values are too low in the winter to relate peaks to climate data directly.

#### 3.2.2 Correlative Statistics

Correlations were sought between monthly values for each climate variable and disease. Using the entire dataset (as opposed to peak-to-peak) allows identification of more subtle relationships than the peak-to-peak analysis, whilst mitigating the impact of fluctuation between different months or seasons.

Using Spearman’s rank analysis, Karnataka HS data negatively correlate with maximum temperature, a medium negative correlation with temperature and DTR, and a medium positive relationship with precipitation. There is also a weakly positive relationship with HS and vapour pressure (Fig. 5).

**Figure 5:**
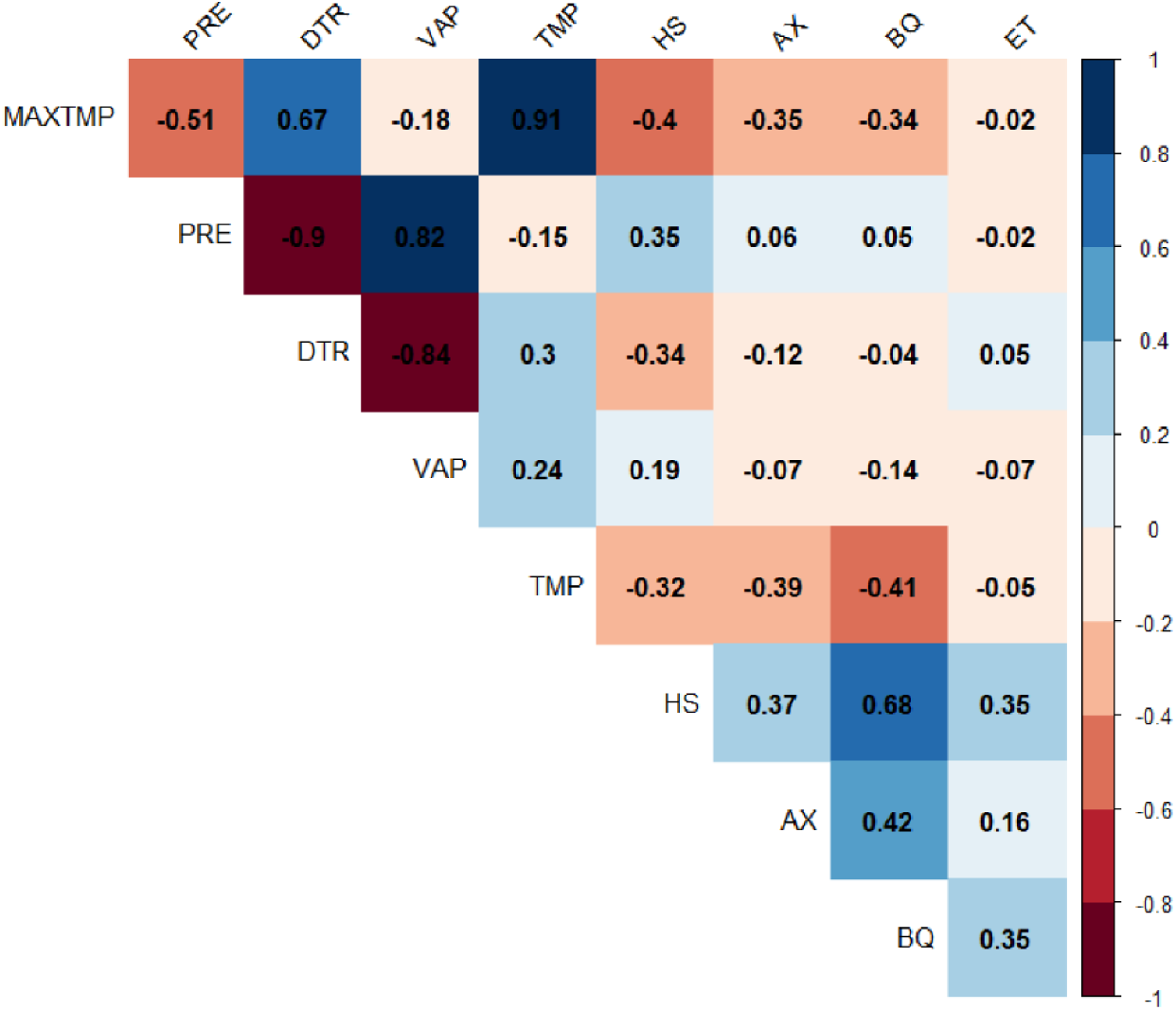
Spearman’s rank correlation plots for climate data vs. bacterial disease. All monthly values over 1987– 2020 are used.

Anthrax has a modest negative relationship with temperature and maximum temperature and a weak negative relationship with DTR. Black Quarter has a negative relationship with maximum temperature and temperature, and no clear relationship with DTR, precipitation or vapour pressure. Finally, ET does not appear to correlate with any of the climate variables.

#### 3.2.3 Principal Component Analysis

Since the statistical correlative tests indicate possible relationships between the disease and climate data, further analysis is appropriate to reduce the dimensionality of the data and define any relationships. This is especially needed as correlative statistics like those in Fig. 4 may not reflect real relationships, only hinting at possible ones. A principal component analysis (PCA) attempts to group the different variables by reducing the overall dimensionality of the dataset and maximising the variance, allowing clearer identification of data relationships and variable contributions to these (Abdi and Williams, 2010). The success of this technique is limited by the quality of the disease data (i.e., accuracy and timespan); however, inferences can still be made regarding relationships between the climate and disease variables.

Three principal components were determined to account for the majority of variance in the datasets. Principal components 1, 2 and 3 account for 81% of the overall data.

Principal Component 1 (Fig. 6A) represents 39.9% of the overall data, predominantly contributed to by maximum temperature (20%), DTR (19%), precipitation (16%) and HS (12.5%). On the other hand, PC2 is primarily contributed to by vapour pressure (25%), BQ (15.5%), precipitation (13%) and DTR (10%). Precipitation and vapour pressure strongly do not depend on DTR, indicating a strong negative relationship. Maximum temperature and temperature are directly opposite to HS, BQ and AX whilst somewhat less, but still opposite, ET, indicating a relatively strong negative relationship between these diseases and climate variables (Fig. 6A). Finally, PC3 is contributed to most by ET (32%), then temperature (24%), maximum temperature (16%) and HS (15%). There is a more notable variation of ET away from the other diseases whilst precipitation, vapour pressure, HS, AX and BQ group together as they have similar PC1 values (Fig. 6B).

**Figure 6:**
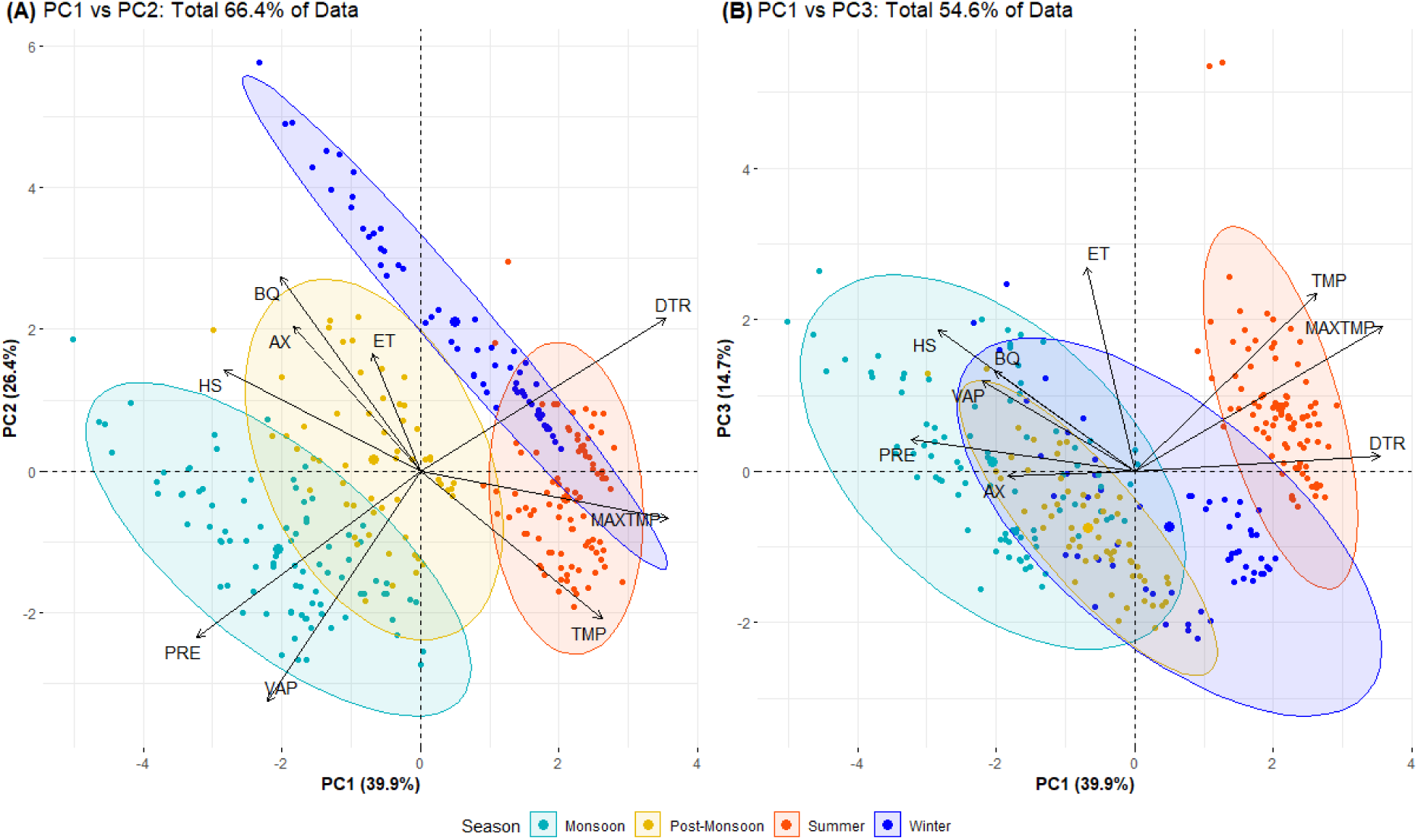
Principal component analyses for Karnataka, indicating relationships between climate variables and bacterial disease data. (A) PC1-PC2; (B) PC1-PC3. Data per season are highlighted by the colour groups.

Based on these groupings, PC1 could be a representation of relative climate moisture, with higher values associated with drier periods (summer and winter data grouped more positively) whilst lower values reflect wetter periods (monsoon data grouped in negative PC1 field (Fig. 6A). Following this, PC2 may represent relative temperature as lower values are mostly the monsoon and summer periods (typically hotter) and then progressively higher values are the post-monsoon then winter periods (typically cooler) (Fig. 6 A). When considering just PCs 1-2, there is no clear relationship between precipitation, vapour pressure and the bacterial diseases; however, they do have similar PC1 eigenvalues, and they become more closely grouped when considering PC1-3 with the exception of ET, which is much more isolated and dominates the PC3 axis. It can therefore be inferred that PC3 may mostly represent the values for ET, rather than a clear seasonal shift as represented by PC1 and PC2. Grouping of HS, AX, BQ, vapour pressure and precipitation (Fig 6B) indicates a positive relationship between these variables within at least 54.6% of the data. The mean values for HS, BQ and AX all fall within the monsoon, winter, and post-monsoon ellipses, indicating that these seasons are the most dominated by bacterial disease outbreaks compared to the Indian summer period.

### 3.3 Predicting Bacterial Disease Outbreaks

The identification of relationships between bacterial disease and specific climate variables is essential for assessing future disease-related risk to livestock potentially affected. Using the defined relationships from Section 3.2, it is possible to define levels of risk for these diseases and project this onto a map of Karnataka to facilitate more spatial interpretation. Even though the disease data collected are at state-level accuracy, we use the defined relationships to project risk at a more granular level, to the resolution provided by the CRU climate data with 0.5 x 0.5° grid boxes.

#### 3.3.1 Present-Day Risk Scenarios

As climate variables can vary significantly throughout the year, it is most important not only to consider the total average for each grid box across Karnataka, but also to focus on the deviation from the mean within each key season that links to the diseases.

When considering the mean of the total climate data for each gridbox, the southern region of Karnataka is prone to higher vapour pressure (23 – 26 hPa) and precipitation (15 – 200 mm) which may indicate a potentially higher risk area to HS, AX and BQ. This is especially relevent when combined with cooler temperature data in the south (24–25°C) and lower maximum temperatures (28–30^°^C). It is however important to consider how different areas of the state may deviate away from these normal values throughout the average year. Areas that may have lower total mean values may still be higher risk if they frequently deviate more so than others. For example, the northern region of Karnataka has a lower vapour pressure average across the period, but in the monsoon and winter seasons it increases significantly (>5 hPa), which could then provide more possibility for disease when combined with higher rainfall and lower temperatures (Fig. 7). This pattern is especially pronounced along the western coastline of Karnataka, where throughout the winter and monsoon, vapour pressure increases with rainfall and a decrease in both temperature and maximum temperature: optimum conditions for HS, AX and BQ.

**Figure 7:**
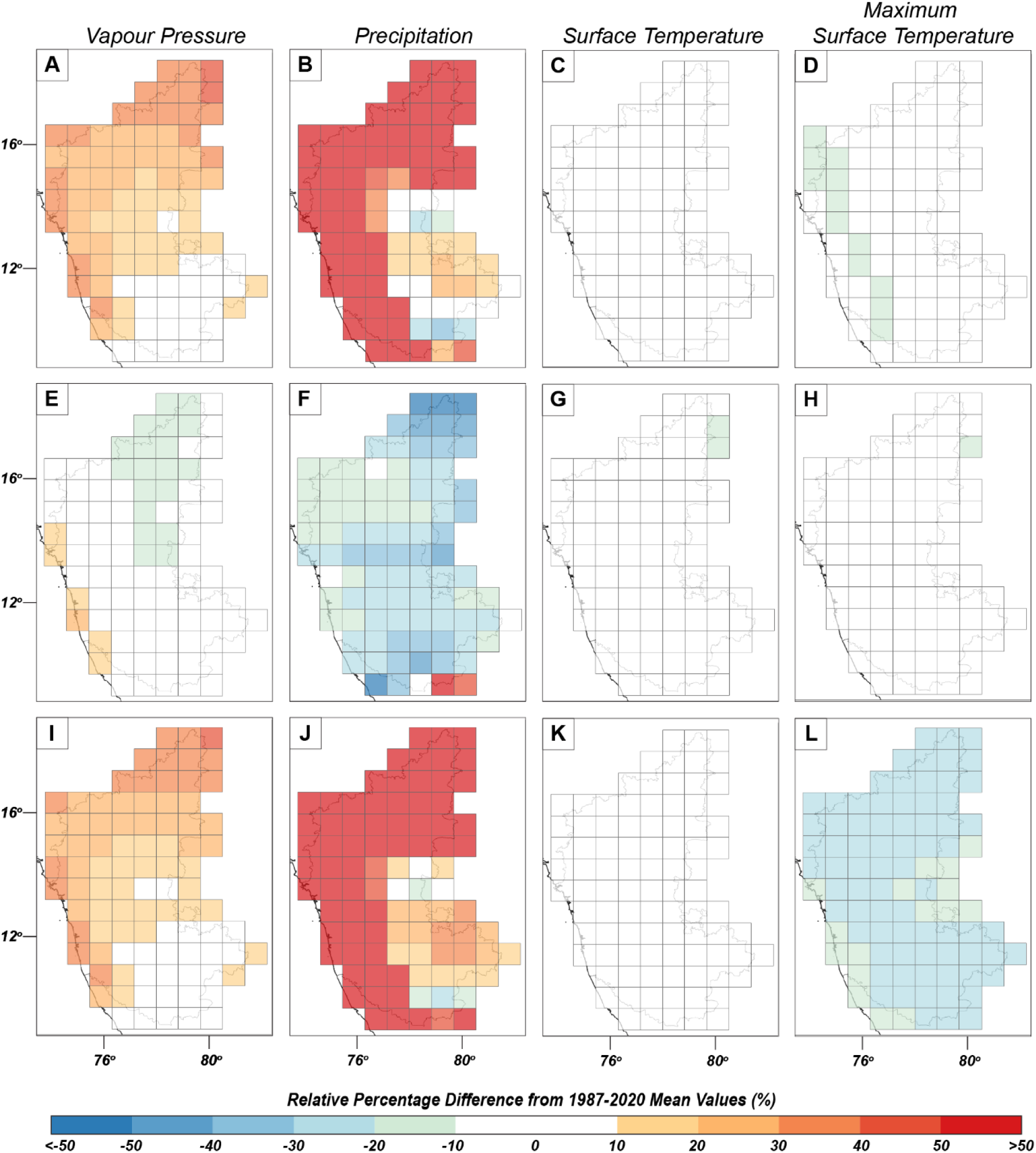
Climate variability maps for the state of Karnataka, showing the average deviation from 1987–2020 mean per climate variable: (A-D) Monsoon; (E-H) Post-Monsoon; (I-L) Winter. Variables are represented by relative percentage difference (RPD, %) for comparison purposes.

When considering risk using climate, it is necessary to consider annual change rather than relying on long-term trends. Throughout the different seasons, different regions of Karnataka are affected by climate variables in different ways, making the assessment of risk hard to define. To mitigate this, risk needs to be established per season and then combined into a total assessment so that it can be looked at with as much temporal variability in the data already accounted for.

When the impact of each climate variable is combined and assessed as one overall factor, it is possible to assign each grid box with a level of risk. The level of risk here has been given a traffic-light system from ‘very low risk’ (dark blue) to ‘very high risk’ (dark red) (Cavan and Kingston, 2012; Murnane, Simpson and Jongman, 2016; Sahoo and Sreeja, 2017)

This classification system was conducted on individual season data (Fig. 8A–C) to infer how each season affects the risk to different regions and to what lengths. During the monsoon, the highest-risk areas are the north-western coastline and the southwestern area, with relatively lower risk zones more in the central-eastern areas (Fig. 8A). This risk lowers during the post-monsoon period, but there are still multiple zones at high risk (Fig. 8B). The eastern and southern areas have increased risk during the post-monsoon season. Lastly, winter poses very high risk in the north and north-western regions, with medium-low risk areas concentrated more in the central-eastern and south-eastern regions (Fig. 8C)

**Figure 8:**
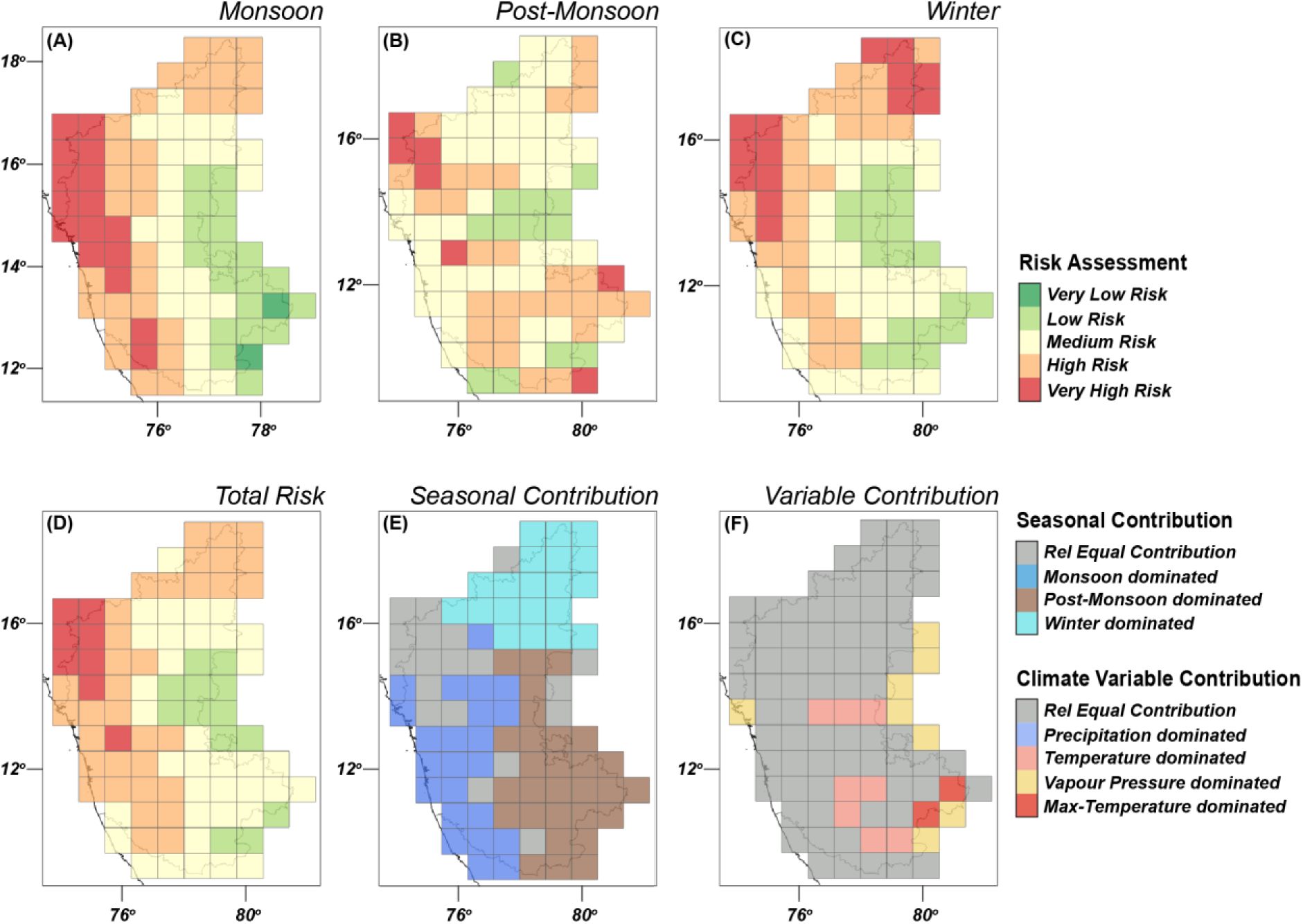
(A-D) Seasonal and overall risk maps based on climate variables. (E) Contribution of each season to the overall risk projected in (A). (F) Contribution of each climate variable to the overall risk projected in (A).

Although useful for identifying seasonal variation in risk, these maps alone are not the best interpretation of risk. The total risk for each grid box needs to be calculated by aggregating the risk per season into one output. This was again classified using a traffic-light system based on percentiles of the data. It was then possible to visualise areas of low risk and high risk annually, providing a clearer picture for farmers and policy makers (Fig. 8D).

Overall, the north-western coastline of Karnataka is the highest risk area for the bacterial diseases HS, BQ and AX (Fig. 8D). The northern edge and western region of Karnataka are at high risk with the central area remaining at medium-low risk overall. The lowest risk areas are those that remain low during all three seasons mentioned, primarily in the eastern-central region and a few areas to the south-east around Bengaluru. The highest risk zones are primarily contributed to by all three seasons, whilst the northern region is more impacted by the winter (Fig. 8E). The monsoon season contributes the most to the high-risk areas of the western coast and south-western zones and then finally, the post-monsoon dominates contribution to the eastern and south-eastern areas risk. The majority of the map are contributed to relatively equally by each climate variable (Fig. 8: F), although temperature and vapour pressure play more of a significant role in certain grid boxes, typically in the eastern and south-eastern zones.

#### 3.3.2 Future Risk Scenarios

It is possible to extend these risk maps into the future using Coupled Model Intercomparison Project Phase 6 (CMIP6) modelled climate predictions (Eyring *et al*., 2016). Over 2040–2069 it is predicted that SW Indian surface temperatures will increase between 1.7–3^°^C (RC4.5 and RCP8.5 models) and precipitation will increase by 10–30% (Mishra and Lilhare, 2016; Bisht *et al*., 2019). Models also indicate an increase in vapour pressure, although obtaining accurate measurements is difficult.

Several scenarios have been modelled using the CRU 4.5 precipitation, temperature and vapour pressure data used in this study, and adjusting the data to these future possible values (Harris *et al*., 2020).

In scenario 1, the seasonal RPD from the 1987–2020 mean was adjusted by adding 2°C to each temperature and maximum temperature value, 10% of itself to the precipitation value (Kulkarni *et al*., 2020; Sanjay *et al*., 2020), and 0.8 hPa to the vapour pressure value (Shrestha *et al*., 2020). This was repeated in scenarios 2 and 3, but instead using 20% then 30% rainfall adjustments, and 1 hPa then 1.2 hPa vapour pressure adjustments. The 2^°^C temperature adjustment was kept the same as this seems the most confident prediction from CMIP6 models, while rainfall and vapour pressure are more uncertain. Future precipitation and vapour pressure changes were chosen based on long-term historic trends, CMIP6 modelled values and literature that fits within realistic parameters (Chattopadhyay and Hulme, 1997; Sun *et al*., 2018; Ha *et al*., 2020; Schwingshackl *et al*., 2021). Risk classifications were then assigned in the same manner as the present-day risk maps.

Using these scenario changes in temperature, precipitation, and vapour pressure there is no notable change in risk from the present-day through to the longer-term future (>40 years); therefore the ‘Total Risk’ panel (Fig 8D) is an accurate representation of present-day risk and future risk (within these models). The north-western area remains highest risk, and the northern and southwestern areas are at high risk. In scenarios 2 and 3, there is a change in one grid box near Bengaluru, to the southwest, from medium to low risk; however, that is the only change, despite a modelled increase in precipitation and vapour pressure. Future work to integrate more accurate climate modelling should enhance the accuracy of these maps, involving iterative updates with additional time series.

## 4 Discussion

### 4.1 Bacterial Disease Trends and Future Predictions

Indian farmers and veterinarians are already aware of the impacts of climate change on their livestock and livelihoods and concerned about what the future holds (Cole and Desphande, 2019; Greru *et al*., 2022). New tools are required that will enable them to better predict and mitigate those changes or give them confidence that their livelihoods will not face increased challenges in the short-to-mid-term future. Our use of correlative statistics and PCA has proven effective in identifying relationships between climate variables and the bacterial diseases HS, AX and BQ. Using peak-to-peak correlations alone cannot identify accurate relationships (e.g., in Fig. 4, it appears the climate variables can both positively and negatively correlate with disease, depending on the year). The main use of the peak-to-peak correlations is to identify certain anomalous spikes in disease and climate trends, such as summer 2008 where ET outbreaks appear to spike significantly (>150). Although the overall trend of HS and BQ appear decrease from the long-term running average (Fig. 3), this may be the result of targeted vaccination programmes instigated by Government of India agencies such as the Department of Animal Husbandry and Dairying (Rathod, Chander and Bangar, 2016; Basu, 2020). These programmes seem to be working (Kushram *et al*., 2020), but could be even more useful if targeted to areas that have been identified at higher risk using the climate data.

From the PCA it has been identified that both vapour pressure and precipitation have a positive relationship with HS, AX and BQ. We showed that these climate variables do not have a relationship with outbreaks of ET. Climate and seasonality have a distinct relationship with HS, AX and BQ, which is in agreement with other studies (e.g. Bhattacharya *et al*., 2005; Bisht *et al*., 2006; Sivakumar, Thennarasu and Rajikumar, 2012).

The preferential seasons for outbreaks of HS, AX and BQ seem to be the monsoon, post-monsoon, and winter periods. This is logical when considering the likely climate variable contributions within each season, and the ideal parameters for outbreak (higher precipitation and vapour pressure, lower temperatures). During the monsoon season, these optimal conditions are met, especially along the Karnataka coastline which is what leads to this area being the highest risk during the June-September period. During the post-monsoon season, there is much lower precipitation; however, vapour pressure remains relatively high along the coast. Temperatures are, however, close to the period average, leading to a decline in risk level across the western region. Winter variable deviation is rather similar to those in the monsoon, with higher precipitation and vapour pressure to the western and northern regions, and cooler maximum / average surface temperatures, leading to increased risk again in the west coast and the highest risk to be in the north. The variation of risk between seasons is critical to defining action taken by farmers and government to mitigate the potential for disease outbreaks all year round.

It is also important to note the way in which risk fluctuates into the long-term future, as establishing new farms or businesses in regions that may increase in risk is impractical. According to our scenario models, there is no significant change in climate-related risk across Karnataka in the long-term future (i.e., 2040–69), a good indication of stability of both local and state-level economies. Increasing rates of climate change may still affect this, however, and consistent updating with the latest climate predictions should be conducted to ensure risk assessment remains as accurate as possible. One further consideration is the level of expertise in extracting and analysing data that is currently required to use our models, which local farmers are unlikely to meet; co-automating the process with local partners is therefore an important next stage of the process.

### 4.2 Limitations of Bacterial Disease Data - NADRES v2

Success criteria rely on the quality of both climate and disease data being used. The NADRES v2 database was the source of disease outbreak data; although it proved useful for studying Karnataka, the data do not seem as robust for all regions of India. For example, disease data for the state of Assam are limited to the year 2000 onwards, and even within this period data seems full of artefacts resulting from the original collection process (e.g., anomalous spikes then years of no data). Data are also restricted to outbreaks alone; no data are available for exact numbers of cases or deaths; assumptions had to be made against general trends and assumptions that farmers reported uniformly across all years, so that a year in which a high number of outbreaks were recorded was actually indicative of more outbreaks rather than just more reported ones.

The interface itself is also not the most user-friendly with respect to data extraction, as automatically generated graphs are missing the axis values or are limited to set parameters. Manual collection is therefore the only way to extract data, using the GIS interface, which itself has numerous usability issues. The number of diseases with data available is limited and the long-term range of the data is restricted to earliest in 1987, making longer-term studies impossible without other data sources. More open-source data with improved user access is critical to future investigations using livestock disease data in India. Although the diurnal temperature range data was not useable for Karnataka, the overall CRU TS 4.5 series of data proved excellent in both quality and accessibility. Epidemiological data matching the granularity and timespan of the CRU dataset would be ideal for future data-integrated investigations, and would address noted concerns with the current lack of standardised data over long-term (i.e. decades, centuries and even millennia) available for rigorous testing of socioecological system resilience, making such long-term predictions difficult (Allen *et al*., 2014).

### 4.3 Considerations for Application of Risk Maps

The risk maps generated from this study use climate variables alone as a classifying parameter. Risk calculations were based on relative percent differences of seasonal averages from total period (1987–2020) averages. Classification of risk then uses sequential 0.2 percentile ranges of the RPD values to assign a risk category to the specific grid box. This system is therefore sensitive to RPD values, which may not indicate a significant change from the period mean or are all either negative or positive. Using this system requires user interpretation of the initial RPD values and raw data to ensure risk assignment is objective. For example, our surface temperature RPD values are lower than precipitation RPD values, as rainfall fluctuates more than temperature. These values require user verification of real temperature anomalies, and that the percentile ranges reflect the increase/decrease in deviation. Without manually checking, it is possible that the ‘very high risk’ category using the 0.8 percentile could match with a negative number instead of values >0. This risk classification system also gives no preferential weighting to any particular climate variable. Future work should be geared towards defining relationships between climate and these diseases. Once clearer thresholds are established, preferential weightings could be applied to the risk assignment e.g., if vapour pressure were to have a significantly greater impact on HS outbreak than temperature, it should contribute a higher weighting to risk classification.

### 4.4 Future Planning and Policies for Poultry / Livestock Farms

Mitigating the impact of climate on livestock is a long-term problem and therefore requires advanced planning with effective long-term solutions. Our recommendations are three-fold. First, future farming and livestock policies need to be implemented that fully respect the longevity of the impact of climate on disease outbreak and mitigate this effectively. Secondly, farmers themselves need to be better informed and able to make local decisions to address the particular variable(s) that may impact them the most. Thirdly, future research should continue to further define these meteorological-epidemiological relationships and classify distinct thresholds further.

Our recommendations address and acknowledge the quality of disease data collection. Available disease data via NADRES have insufficiently high spatial resolution; however, we have identified parameters that may relate to increased outbreak risk in certain bacterial diseases on a state level (i.e., precipitation, vapour pressure). By disaggregating these parameters to the resolution of the climate data (0.5 x 0.5°), risk mapping can be conducted at a higher resolution than the original disease data provided. However, if disease data were to be made more easily available at a much more granular level and frequency, improved interpretations, and accuracy of the levels of risk would result.

Typical risk assessments follow hazard-orientated procedures; similarly, we identify and model these complex relationships and define critical relationships. A definition of critical thresholds per disease and per potentially impacted livestock, however, would be far more beneficial. This risk assessment provides an insight into larger regions and into long-term planning more than specifically providing disease critical thresholds per climate variable. Further investigations should be carried out to define quantitative thresholds for precipitation and vapour pressure to which disease outbreak is related.

One possible solution to mitigating the impact of climate change on livestock is the wider introduction of environmentally controlled sheds (Ambazamkandi *et al*., 2015), along with the energy infrastructure needed to support them; such infrastructure is currently insufficient in many rural and even peri-urban areas (Greru *et al*., 2022). A second consideration is a shift from livestock and poultry rearing to aquaculture, as is already being seen in northeast India, in regions likely to become more prone to heavy rainfall and flooding (Sarkhel, 2015; Rao, 2017); or shifting farming operations to different areas. Further solutions include the introduction of other technologies e.g. vertical farming to help sustain crop farming (Benke and Tomkins, 2017; Maheshwari, 2021), especially in particularly challenging regions where climate stress challenges both crop and livestock operations. Establishing future livestock farms in areas of consistent low risk is feasible, as well as the modification of existing farms, which should be made relative to the highest contributing risk-causing variable. For instance, support should be directed to areas where we define the monsoon season as the highest risk (due to the influences of precipitation and vapour pressure) to address the impact of saturated soil, wet feed, and humid living conditions on livestock (Pathak, Aggarwal and Singh, 2012).

## 5 Conclusions

We have identified a clear awareness of climate change impacts amongst those whose livelihoods depend on farming and have developed a system through which risk can be better understood and predicted. Our efforts evidence a relationship between average and maximum surface temperature, precipitation and vapour pressure, and several livestock bacterial diseases. There is a modest positive relationship between precipitation and vapour pressure with HS, AX and BQ, followed by a negative relationship between temperature and maximum temperature with the same diseases over a period of 30 years. There is no identified relationship between ET and diurnal temperature range and these climate variables.

Based on these relationships, we find that the north-western coast of Karnataka is the highest-risk area for HS, AX and BQ, irrespective of other factors that may also govern outbreaks. The western coastline and northern regions are at high risk of outbreak, while the central-eastern and south-eastern regions are the lowest risk. These risk levels are not predicted to change in the next 50 years, even with increased temperatures, and changing spatiotemporal patterns of precipitation and vapour pressures following CMIP6 modelled values. This may not, however, be true of other regions of India, or globally, where changing climate conditions over the coming decades are likely to shift the climate parameters of currently low-risk regions into higher-risk states. This suggests that the ability to predict climate fluctuations and long-term changes will become increasingly important in coming decades and may require greater consideration of climate science within policy intended to protect and improve animal health, such as increased joining up of the UNFCCC and the Global Action Plan on AMR (known colloquially as the ‘Tripartite Agreement’) agreed by the World Health Organization (WHO), Food and Agriculture Organization of the United Nations (FAO) and the World Organization for Animal Health (OIE) This will be particularly in regions such as India that are at the sharp edge of that change (Rajesh, 2021). In short, we argue that animal health cannot be considered independently of climate change. Such considerations may also help to converge fields that approach challenges to global health from slightly different angles, such as One Health and Planetary Health, and offers to unite them behind common goals.

Epidemiological data and interpretations were restricted to Karnataka; data for other states of interest (i.e., Assam, where NEOSTAR was conducted) were too limited in both time and space to provide insight. This is due to the poor data availability through the NADRES vs database. Our work could add to the NADRES online system by providing long-term predictor maps for India livestock disease; the existing maps provide only two-months’ notice of increased risk.

The techniques used here can be applied to analogous projects for multi-purpose use, particularly for those where climate and epidemiological data cover matching time frames at equal resolution (e.g., monthly averages). The use of this workflow to generate long-term risk maps can also be applied elsewhere in the world. Our future intentions are to automate the risk map production process and then test the model on epidemiological datasets that are equally robust and granular as the climate data.

## Acknowledgements

For open access purposes, we have applied a Creative Commons Attribution (CC BY) licence to any Author Accepted Manuscript version arising from this submission.”

We also thank Jayant Deshphande, independent poultry consultant to the DARPI project, and Gowthaman Vasudevan, Assistant Professor at Tamil Nadu Veterinary and Animal Science University.

## Data Availability Statement

All meteorological data involved in this study were taken from the CRU 4.5 TS gridded dataset, hosted by CEDA. All epidemiological data used were collected from the NADRES v2 GIS platform hosted by NIVEDI. Both databases are online and publicly available (ICAR-NIVEDI, 2017; Centre for Environmental Analysis (CEDA), 2022).

